# *Vibrio cholerae* pathogenicity island 2 encodes two distinct types of restriction systems

**DOI:** 10.1101/2024.04.04.588119

**Authors:** Grazia Vizzarro, Alexandre Lemopoulos, David William Adams, Melanie Blokesch

## Abstract

In response to predation by bacteriophages and invasion by other mobile genetic elements such as plasmids, bacteria have evolved specialised defence systems that are often clustered together on genomic islands. The O1 El Tor strains of *Vibrio cholerae* responsible for the ongoing seventh cholera pandemic (7PET) contain a characteristic set of genomic islands involved in host colonisation and disease, many of which contain defence systems. Notably, *Vibrio* pathogenicity island 2 contains several characterised defence systems as well as a putative Type I restriction-modification system (T1RM), which, interestingly, is interrupted by two genes of unknown function. Here, we demonstrate that the T1RM system is active, methylates the host genomes of a representative set of 7PET strains, and identify a specific recognition sequence that targets non-methylated plasmids for restriction. We go on to show that the two genes embedded within the T1RM system encode a novel two-protein modification-dependent restriction system related to the GmrSD family of Type IV restriction enzymes. Indeed, we show that this system has potent anti-phage activity against diverse members of the *Tevenvirinae*, a subfamily of bacteriophages with hypermodified genomes. Taken together these results expand our understanding of how this highly conserved genomic island contributes to the defence of pandemic *V. cholerae* against foreign DNA.

## IMPORTANCE

Bacterial defence systems are specialised immunity systems that allow bacteria to counter the threat posed by bacterial viruses (bacteriophages) and other invasive mobile genetic elements such as plasmids. Although these systems are numerous and highly diverse, the most common types are restriction enzymes that can specifically recognise and degrade non-self DNA. In this work we show that the *Vibrio* pathogenicity island 2, present in the human pathogen *Vibrio cholerae*, encodes two types of restriction systems that use distinct mechanisms to sense non-self DNA. The first is a classical Type I restriction-modification system that recognises specific DNA sequences, which are protected in the host genome by methylation. The second, is a novel modification-dependent Type IV restriction system that recognises hypermodified cytosines present in certain bacteriophage genomes. Curiously, these systems are embedded one within the other, forming a single cluster, suggesting that the systems collaborate to create a multi-layered defence system.

## Introduction

Mobile genetic elements (MGEs) such as plasmids, transposons and integrative-conjugative elements can confer significant fitness advantages by facilitating the transfer of beneficial traits to the host bacterium, including key virulence factors or antibiotic-resistant genes (1). However, their maintenance and or activity can also impose a metabolic burden on the host cell, while elements that integrate on the chromosome have the potential to disrupt important genomic features (2). Furthermore, the replication of some MGEs results in the death of the host cell (3). Indeed, predation by lytic bacteriophages (phages), which are ubiquitous bacterial viruses, has imposed a strong evolutionary pressure to develop multiple lines of defence against these elements, including a vast array of specialised defence systems (4, 5).

Upon recognising an infection, these systems can either respond directly by degrading the invading non-self-DNA and thus provide individual level protection, or alternatively can sacrifice the host cell prior to phage induced lysis to protect the surrounding population (abortive infection, Abi) (6). The most common and best-studied defence systems are Restriction-modification (RM) systems, which use restriction enzymes to directly degrade non-self DNA (5). Type I-III RM systems are modification-blocked enzymes that recognise specific DNA sequences and only cut DNA when it is unmodified, while the corresponding sequences in the host genome are protected by epigenetic modification with a cognate methylase (7–9). In contrast, Type IV systems are modification-dependent enzymes that can recognise and degrade invading DNA with specific modifications, which are used by some phages to avoid restriction by modification-blocked systems (9, 10).

Diverse defence systems, including RM systems, tend to cluster together within genomic islands known as “defence islands” (11, 12). This pattern also applies to the defence systems identified so far in *Vibrio cholerae*, the causative agent of cholera. This bacterium features specialized islands crucial to its pathogenic evolution. Indeed, only certain *V. cholerae* strains, referred to as toxigenic isolates, can cause cholera. This ability is due to the presence of two key virulence/colonization factors: the cholera toxin (CT) and toxin-coregulated pilus (TCP), encoded on the filamentous phage CTXΦ and the *Vibrio* pathogenicity island 1 (VPI-1), respectively (13–16). The ongoing seventh cholera pandemic is caused by the O1 El Tor *V. cholerae* lineage (7PET), which uniquely carries the *Vibrio* seventh pandemic islands I and II (VSP-I and VSP-II), characteristic of the 7PET strains (17, 18). These genomic islands are implicated in defence, as they encode for instance CBASS and AvcD systems (VSP-I) and the Lamassu system DdmABC on VSP-II (19–23). Additionally, toxigenic *V. cholerae* strains carry the *Vibrio* pathogenicity island 2 (VPI-2), which is believed to enhance pathogenicity by giving the pathogen a competitive advantage in using sialic acid as a carbon source during gut colonization (24–26). This capability is encoded within the island’s *nan-nag* genomic region (24–26). Moreover, the island houses several genes believed to protect against MGEs, including (i) a predicted Zorya Type I system, a phage defence system identified across a wide range of bacterial genomes and experimentally studied primarily through *Escherichia coli* homologs (27, 28); (ii) the DNA defence module DdmDE that targets and degrades small multicopy plasmids (23); and (iii) a gene cluster/operon predicted to encode a Type I restriction-modification (T1RM) system (24). The presence of both predicted and established defence systems encoded within VPI-2 suggests that it may serve as a genuine defence island.

In this study we set out to characterize the predicted T1RM operon within VPI-2. We show that the T1RM system promotes methylation of the genomes of 7PET *V. cholerae* strains, and identify a specific recognition sequence that can target non-self derived plasmids for restriction. Furthermore, we identify two genes embedded within the T1RM operon that form a novel modification-dependent restriction system related to the GmrSD family of Type IV restriction enzymes, which we term TgvAB. When produced in *E. coli* this system has potent anti-phage activity against phages with hypermodified genomes. Collectively, these findings enhance our understanding of how this highly conserved genomic island contributes to the defence of pandemic *V. cholerae* against foreign DNA.

## Results and Discussion

### *In silico* analysis of VPI-2 and the T1RM cluster

Although VPI-2 was discovered over 20 years ago (24), the genes it carries have not yet been fully characterized. To begin bridging this knowledge gap, we started by re-evaluating the conservation of the island. Consistent with earlier findings (24), this revealed that VPI-2 is highly conserved among a set of 7PET O1 strains isolated between 1975 and 2011. Our analysis confirmed the island’s modular structure as described by Jermyn and Boyd (24, 29), including a predicted Zorya system (27) encoded by genes *VC1761-64* (as per reference strain N16961; (30)), a predicted T1RM system (*VC1765-69*), the DdmDE defence module (*VC1770-71)* (23), the *nan-nag* sialic acid utilisation cluster (*VC1773-1784*), and a region with phage-like properties (*VC1791-1809*) (24) (Fig. 1a). Notably, O139-serogroup strains such as MO10 carry a highly truncated version of VPI-2 that retains only the phage-like region (24, 31) (Fig. 1a).

**Figure 1.**
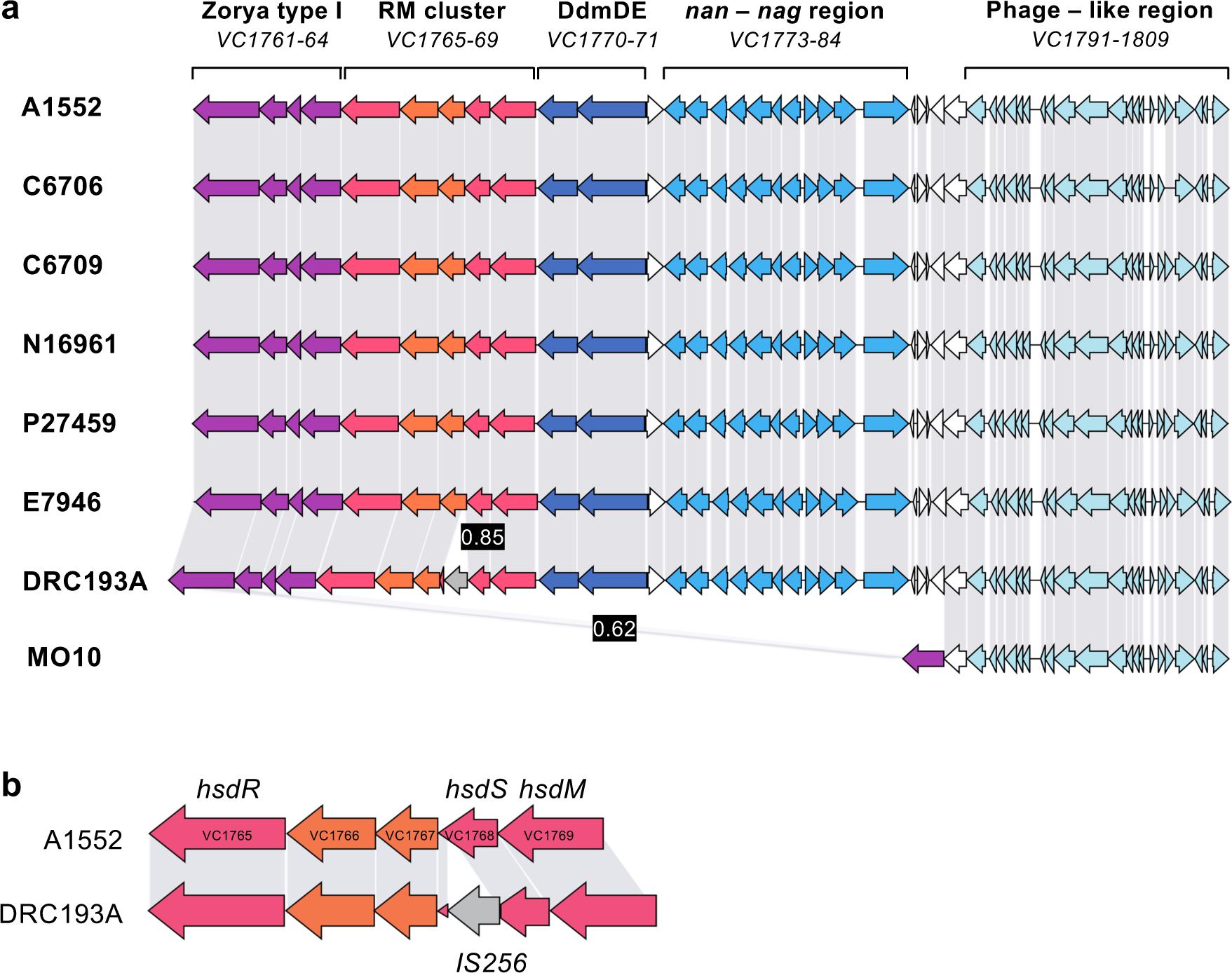
VPI-2 exhibits high conservation across 7PET *V. cholerae* strains. a) Comparative genome alignment of the *Vibrio* pathogenicity island 2 (VPI-2) across a selection of 7PET O1 and O139 strains, isolated between 1975 and 2011. The genomes of these strains are displayed alongside their designated strain name. Coding sequences within the genomes are represented by arrows, with grey bars connecting them to indicate amino acid identity percentages at or above a threshold of 0.93. Instances of lower identity are highlighted in black boxes. Gene locus tags are derived from the reference genome of strain N16961. Predicted or established functions are labelled above each cluster. **b)** Close-up examination of the *VC1765-69* gene cluster in strains A1552 and DRC193A reveals three genes responsible for the components of the putative T1RM system (*hsdR, hsdS, hsdM*). A comparative alignment highlights the disruption of the T1RM cluster in strain DRC193A, caused by an IS*256* transposon insertion within *hsdS*.

Given the observed conservation of VPI-2 and the presence of established defence systems, we explored the possibility that the putative T1RM system was also actively involved in restricting foreign DNA. Interestingly, the previously annotated T1RM region sits within a five-gene operon, of which three genes encode homologs of the known T1RM components (Fig. 1b). These host-specificity determinant (*hsd*) genes encode: the specificity subunit HsdS, which recognises a specific DNA recognition sequence; the methylase subunit HsdM, which methylates (and therefore protects) the recognition sequences in the host genome; and HsdR, the restriction enzyme subunit, which upon encountering foreign DNA with an unmethylated recognition sequence translocates the flanking DNA and cleaves at variable distances from the recognition site (7, 32–34). These components function together as multi-subunit complexes capable of both methylating and restricting DNA. Importantly, restriction requires a pentameric complex of 2HsdR + 2HsdM + HsdS, and although HsdR is dispensable for methylation, HsdS is required for both activities (7, 32). Interestingly, two genes of unknown function are embedded within the T1RM cluster (*VC1766-67*; Fig. 1b), which we characterize in the subsequent sections below.

## Deciphering the recognition motif of VPI-2’s T1RM system

If the T1RM system is active in *V. cholerae* then we predicted that we should be able to detect a specific methylation signature that is absent is strains lacking this system. To test this hypothesis, we used SMRT PacBio whole-genome sequencing, which can detect the presence of various DNA modifications including methylation, to determine the methylomes of a selection of 7PET O1 serogroup strains (strains as in Fig. 1a), as well as those of control strains lacking the T1RM system (see methods) (35–37). As shown in Figure 2a, this analysis revealed a unique 13-nucleotide motif with methylation marks located on the second nucleotide within the sequence GATGNNNNNNCTT (m6A: GATGNNNNNNCTT:2). Upon further examination, we discovered that this DNA motif is present in over 600 copies throughout the genome of each strain and is modified in nearly 100% of cases in all O1 serogroup strains, except DRC193A (Fig. 2a). This phenotype is likely explained by the interruption of *hsdS* in this strain by a IS*256*-like transposase gene (38) (Fig. 1b). Finally, and as expected, the O139 serogroup strain MO10, which is missing the T1RM-encoding region of VPI-2 (Fig. 1a), and both a VPI-2 and a *VC1765-69*-deficient deletion strain (Table S1) all lacked this particular methylation mark (Fig. 2a).

**Figure 2.**
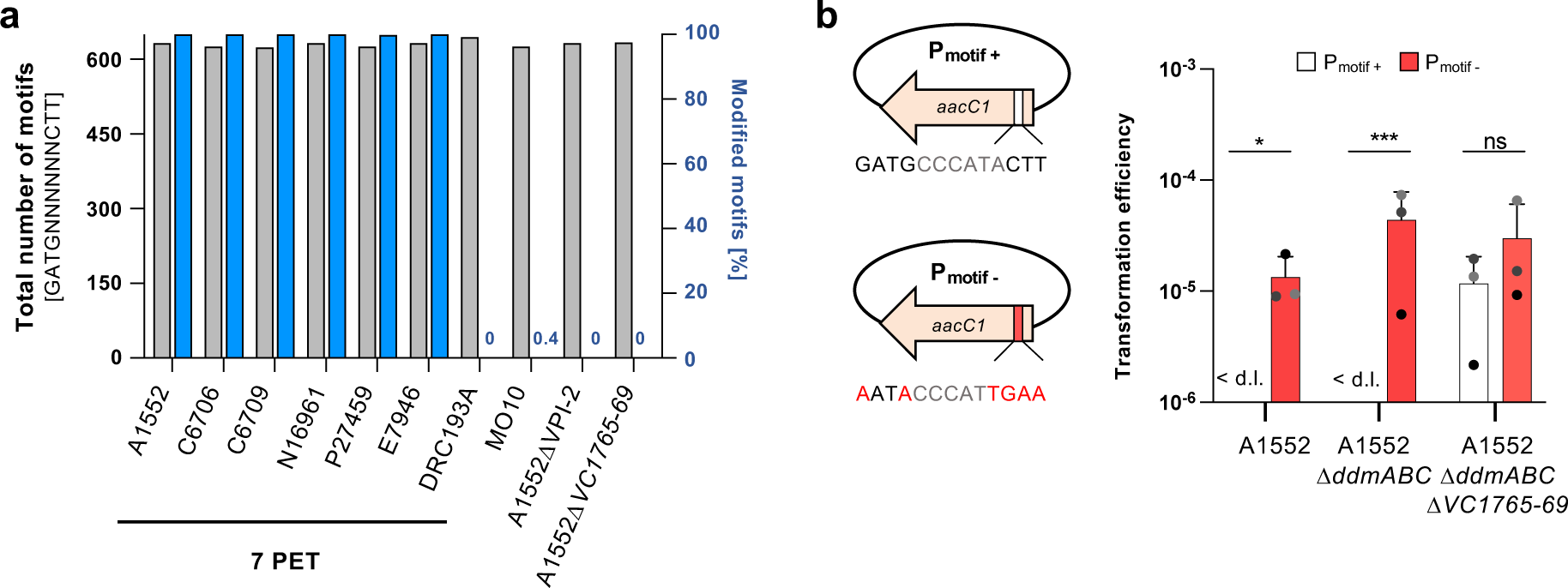
Type I RM system’s role in chromosomal methylation and plasmid restriction. a) SMRT sequencing uncovers a distinctive modified DNA motif across various 7PET *V. cholerae* strains. The grey bars show the number of the DNA motif (GATGNNNNNNCTT) in each genome, while the blue bars denote the percentage of this motif methylated in each strain’s genome (see secondary *Y*-axis on the right). **b)** The T1RM system hinders plasmid acquisition. Transformation assays compare the uptake of a plasmid containing the recognition motif (P_motif+_) against a derivative plasmid with silent mutations altering the nucleotide sequence (P_motif-_). To the left, diagrams of the plasmids are depicted. Statistical differences were calculated on log-transformed data using a two-way ANOVA corrected for multiple comparisons with Šidák’s method. * *P* < 0.05; *** *P* < 0.001; ns, not significant. < d.l., below detection limit.

## The T1RM impairs plasmid acquisition

Having identified the methylated recognition motif, we next tested the ability of this motif to target plasmids for restriction by the VPI-2 T1RM system. Serendipitously, we realised that the recognition sequence is present within the widely used gentamicin resistance cassette *aacC1*. We therefore created plasmid derivatives carrying *aacC1* either with the putative recognition sequence intact (P_motif+_) or with silent mutations that disrupt the nucleotide recognition sequence while preserving the protein coding sequence (P_motif-_) (Fig. 2b). We then purified these plasmids from *E. coli* and used them as substrates in an electroporation-based transformation assay to compare their transformation frequencies in various backgrounds. As shown in Figure 2b, transformation with plasmid P_motif+_ was below the detection limit in the WT background (strain A1552) even though transformants could readily be obtained with plasmid P_motif-_. Furthermore, this disparity between the acquisition of the two plasmids became even stronger in the absence of the DdmABC system (23) (Fig. 2b), which is known to target derivatives of this high-copy number plasmid (39). To determine if the plasmid restriction was mediated by the T1RM system, we removed the corresponding operon from the A1552Δ*ddmABC* background and then assessed the plasmid transformability of the resulting strain. As shown in Figure 2b, deletion of the five-gene restriction cluster on VPI-2 indeed led to the recovery of P_motif+_ transformants. Moreover, the transformation difference between the P_motif+_ and P_motif-_ plasmids was now no longer statistically significant. Consequently, we conclude that the T1RM system is active, that it methylates a specific recognition sequence, and that when this sequence is present on non-self DNA the acquisition of this non-methylated DNA is restricted.

## Genes embedded in the T1RM cluster protect against phages with modified genomes

Type I restriction-modification systems are recognized for their important role in defending the cell against phage infection (40). Therefore, we aimed to investigate the ability of the entire RM cluster, including the two embedded genes, to protect against viral infections. However, given that commonly used *Vibrio* phages, such as ICP1, ICP2, and ICP3, are typically isolated using VPI-2-carrying 7PET strains as the host (for example, strain E7946 and its derivatives (41)), it is unlikely that any defence system encoded on VPI-2 would provide protection against these phages. Therefore, we engineered the *E. coli* strain MG1655 to carry an arabinose-inducible version of the entire five-gene RM cluster (*VC1765-69*), which was integrated into its chromosome. Utilizing this strain and a strain without the cluster as a control, we screened for protection against the BASEL collection, a recently established phage collection that represents the natural diversity of *E. coli* phages (42). As shown in Figure 3a (and Fig. S1 for the data on the entire screen), we noted a reduction in the efficiency of plaquing of at least 1000-fold compared to the non-defence control upon infection with members of the *Tevenvirinae* subfamily. The *Tevenvirinae* subfamily is characterized by their unique cytosine modifications, which play a crucial role in their defence against RM systems like the T1RM (42). Specifically, *Tequatrovirus* group phages feature cytosines that are hydroxymethyl-glucosylated, while *Mosigviruses* possess cytosines that are hydroxymethyl-arabinosylated (10, 43).

**Figure 3.**
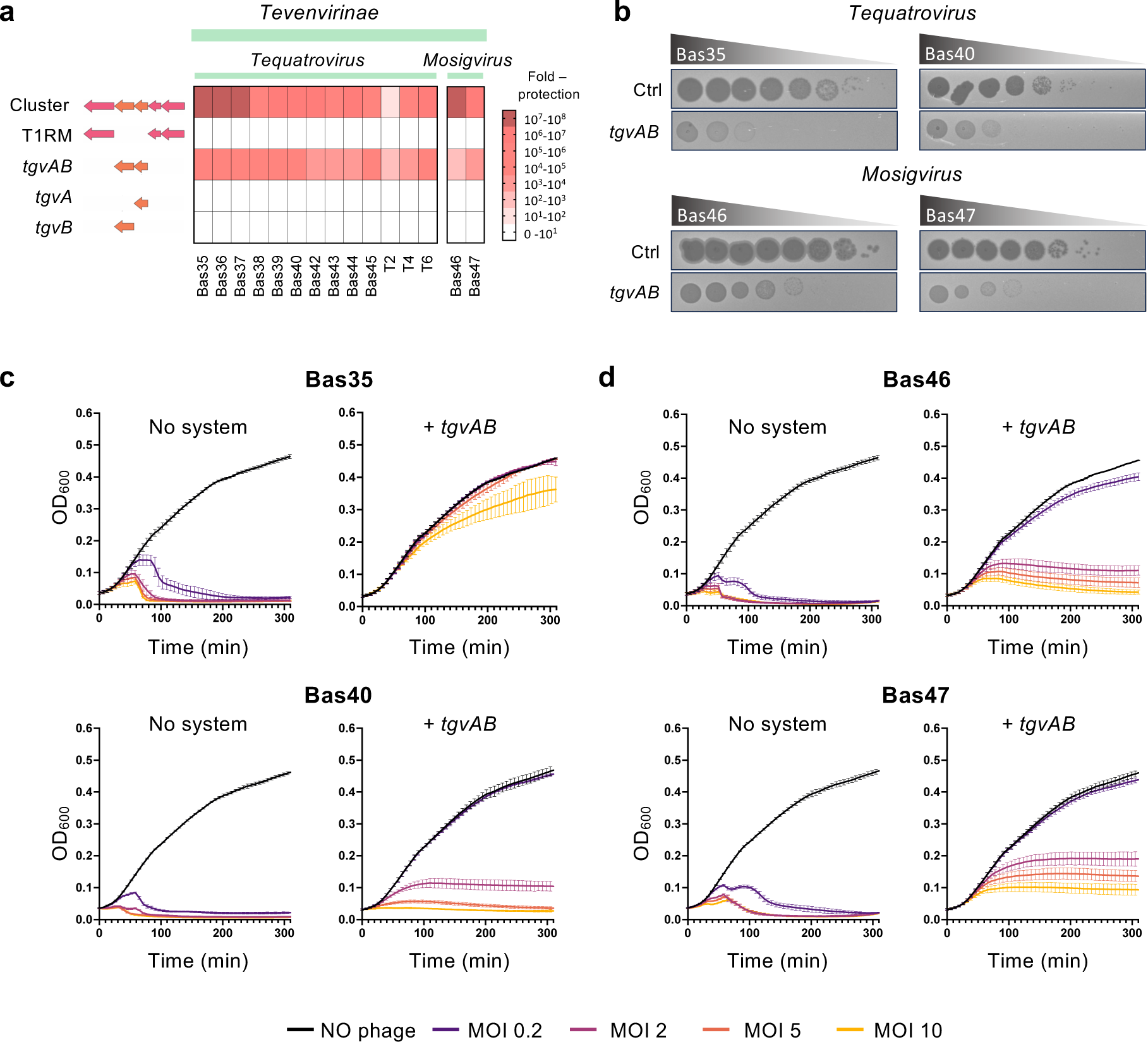
Protection against *Tevenvirinae* by Tgv proteins encoded by the T1RM-embedded genes. a) Observed defence activity against then BASEL phage collection. Protection levels (fold-protection, as shown by the colour code on the right) were determined by comparing plaque formation in strains with the system to those without, using tenfold serial dilution assays. On the left, gene organization of the tested strains. Data represent the average of two replicates. The collective results of all 77 phage infections are detailed in Fig. S1. **b)** Phage plaque assays on *E. coli* strains harboring an empty transposon (control, Ctrl) or the two T1RM-embedded genes (*tgvAB*), using a tenfold serial dilution. **c, d)** Growth curves of *E. coli* cultures carrying an empty transposon (no system) or Tn*tgvAB* (+ *tgvAB*), without (NO phage) or with exposure to phages, initiated at time 0 with various MOIs (0.2, 2, 5, or 10). **c)** *Tequatroviruses* and **d)** *Mosigviruses* were used for infection. The presented data are the average of three independent experiments (±SD, illustrated with error bars).

To determine which part of the RM cluster was responsible for this protection, we created *E. coli* strains that either carried the T1RM cluster or just the embedded two-gene cluster independently. Strikingly, this revealed that the two-gene cluster alone was responsible for this protection (Fig. 3a and b). Furthermore, the two genes did not provide protection when expressed individually, indicating a necessity for their combined action to achieve the observed anti-phage activity (Fig. 3a). For reasons explained below, we named these two genes as **<u>T</u>**ype I-embedded **<u>G</u>**mrSD-like system of **<u>V</u>**PI-2, *tgvA* (*VC1767*) & *tgvB* (*VC1766*).

To dissect the underlying mechanism of anti-phage defence by TgvAB we monitored the growth kinetics of *E. coli* strains infected with increasing multiplicities of infection (MOI) for both *Tequatrovirus* (Fig. 3c) and *Mosigvirus* phages (Fig. 3d). As expected, cultures of the no system control strain grew and then lysed in an MOI-dependent manner (Fig. 3c-d). In contrast, TgvAB producing cultures infected with the *Tequatrovirus* Bas35 continued to grow at rates indistinguishable from those of the no phage control up to and including MOI 5, before being partially overcome at MOI 10 (Fig. 3c). This phenotype is consistent with TgvAB acting directly to target the invading phage. However, TgvAB producing cultures infected with either the *Tequatrovirus* Bas40 or the *Mosigviruses* Bas46 and 47 all showed more variable levels of protection (Fig. 3c-d). Indeed, while protection was robust at MOI 0.2, at higher MOIs we observed growth inhibition and even partial lysis. Nevertheless, given that the cultures mostly continued to grow past the point at which they lysed in the no system control, together with the direct protection observed against Bas35 at all tested MOIs, we conclude that TgvAB likely also acts directly against these phages, but that they are better able to overwhelm the system at high MOI.

## The TgvAB defence system is a member of the GmrSD family of Type IV restriction enzymes

Bioinformatic analysis of the TgvAB system revealed that TgvB (*VC1766*) possesses two domains of unknown function (DUF), an N-terminal DUF262 domain and a C-terminal DUF1524 domain. In contrast, TgvA (*VC1767*) is predicted to carry only an N-terminal DUF262 domain (Fig. 4a-b). Interestingly, previous work by Machnicka *et al.* found that GmrS and GmrD proteins contain the DUF262 and DUF1524 domains, respectively, typically coming together to form GmrSD fusion proteins (44). Notably, the TgvB homolog from classical biotype *V. cholerae* (VC0395_A1364) was also identified as a GmrSD homolog in this study (44). These double domain forms of GmrSD function as modification-dependent Type IV restriction enzymes, and are known to specifically recognise and cleave DNA containing sugar-modified hydroxy-methylcytosine. However, they exhibit no activity against unmodified DNA (44–47). Given that such modifications are typical of the *Tevenvirinae* (10) and the specific protective effect we observed against them (Fig. 3), this suggests that TgvAB may function in a similar manner. Importantly, and in contrast to classical single protein GmrSD such as Eco94GmrSD (Fig. 4a) (45), our phage infection assay revealed that TgvA and TgvB cannot function independently, and that both proteins are required for anti-phage activity.

**Figure 4.**
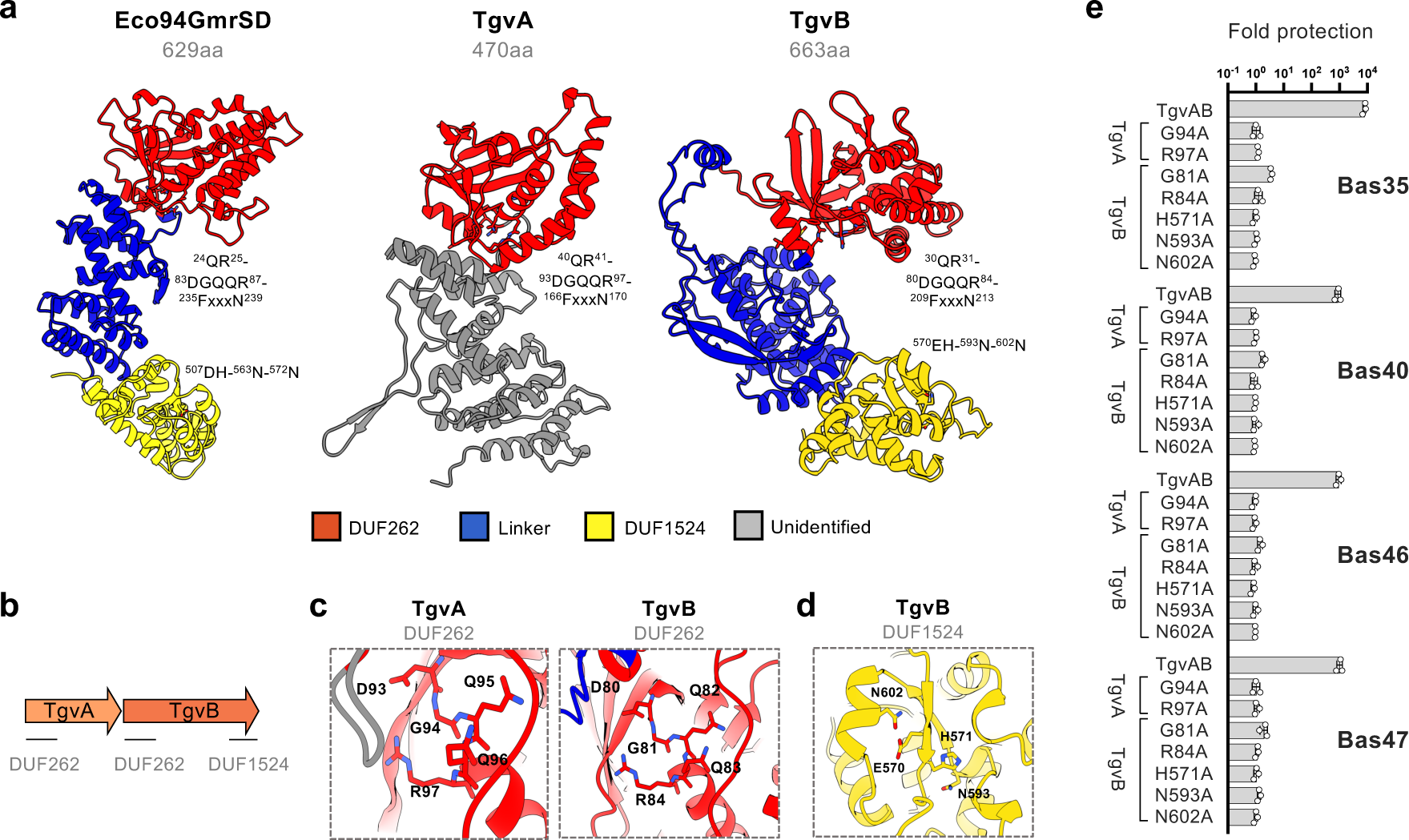
The two-protein TgvAB defence system is a member of the GmrSD family of Type IV restriction enzymes. a) Structural models of Eco94GmrSD of *E. coli* STEC_94C and TgvA (*VC1767*) & TgvB (*VC1766*) of *V. cholerae* 7PET strains. The models, produced via AlphaFold (ColabFold), portray the domains with corresponding colours, while also highlighting the residues characteristic to the DUF262 and DUF1524 domains. Images were generated using ChimeraX 1.7.1 **b)** Schematics displaying conserved domains identified in the TgvA and TgvB proteins. **c, d)** Zoomed view of the conserved (**c**) DGQQR motif found in the DUF262 region of TgvA & TgvB and (**d**) of the His-Me finger motif within the DUF1524 of TgvB, highlighting the catalytic histidine (H) situated at the terminus of the β1 strand, the Asparagine (N) residue positioned in the loop region and the final N residue within the α-helix. **e)** Site-directed mutagenesis removed the antiviral effect. The level of protection was evaluated as described in Fig. 3. Mutagenesis aimed at disrupting NTPase or endonuclease functions exerted by DUF262 and DUF1524, respectively. The data are averages from three independent experiments (±SD, as shown by the error bars).

Machnicka *et al.*, showed that the predominant form of GmrSD is as a single multi-domain protein containing an N-terminal DUF262(GmrS) domain and a C-terminal DUF1524(GmrD) domain, separated by an alpha helical linker region (44). This domain organisation was subsequently confirmed by crystal structures of the related GmrSD family members BrxU, which also recognises and degrades DNA containing modified cytosines, and the phosphorothioate modification sensing enzyme SspE (48–50). Furthermore, biochemical experiments with these enzymes have shown that the N-terminal DUF262 likely functions as DNA modification sensor, and uses nucleotide binding and hydrolysis to regulate the activity of the C-terminal DUF1524, which functions as a nuclease to degrade non-self-DNA (48, 50). Strikingly, structural modelling of Eco94GmrSD and TgvAB using AlphaFold (51), revealed that TgvB is predicted to share a similar domain architecture, although in the case of TgvA, this similarity is limited to the N-terminal DUF262 domain (Fig. 4a-b). Moreover, the top hits in structural alignments of the TgvAB models were SspE and BrxU, reinforcing the idea that these proteins are related.

Next, to further investigate the relative contributions of the DUF262 and DUF1524 domains to TgvAB function, we used the structural modelling and alignments to identify key residues in each domain. For both TgvA and B, the three highly conserved motifs characteristically associated with the DUF262 domain (i.e. (i) QR, (ii) DGQQR and (iii) FxxxN) were readily identifiable (Fig. 4a-d) (44). Notably, the DGQQR motif is thought to form part of a nucleotide binding pocket and to be required for nucleotide hydrolysis. Indeed, site-directed mutants of either TgvA or TgvB encoding substitutions in this motif previously shown to disrupt NTPase activity (48–50), all resulted in a total loss of anti-phage activity (Fig. 4c-e). In contrast, the DUF1524 domain contains a highly conserved H…N…H/N motif, which belongs to the His-Me finger nuclease superfamily and that assumes a characteristic ββα fold (52, 53). Such a motif was readily apparent in C-terminal domain of the predicted TgvB structure, and consistent with previous findings (45, 48–50), substitutions designed to disrupt either the catalytic histidine (TgvB[H571A]) or the metal-binding asparagine (TgvB[N602A]) were sufficient to abolish anti-phage activity (Fig. 4d-e).

Overall, our results suggest that the TgvAB system senses phages with hypermodified cytosines in a manner that requires the DUF262 domains of both TgvA and B, and that the His-Me nuclease domain of TgvB likely functions as the effector against phage DNA. Nevertheless, why TgvB alone is not sufficient for phage protection remains unclear. One possibility is that TgvA is required to overcome a phage encoded inhibitor. For example, some GmrSD family enzymes such as Eco94GmrSD are inhibited by the protein IPI*, which is co-injected into the host cell with the T4 genome (45, 46, 54). However, TgvA could equally also play a regulatory or structural role and further work will therefore be needed to clarify these possibilities.

## Occurrence of the *tgvAB* system within and outside T1RM clusters

To investigate the prevalence of *tgvAB* homologs within the T1RM cluster, we examined the distribution of the specific five-gene operon within 81,172 bacterial genomes (see methods for details). This *in silico* analysis revealed that the gene architecture found in VPI-2 of *V. cholerae* is also present in a variety of other bacterial genera (Fig. 5a) with 102 hits within this genome database, including several *Shewanella, Acinetobacter*, and *Pseudoalteromonas* species (see Table S1 for species-level details). This wider distribution indicates the potential functional conservation of these gene arrangements across different gram-negative bacteria. However, the genus *Vibrio* was still most prominently featured in these findings with 55 hits (Fig. 5a). Precisely, apart from *V. cholerae*, species such as *Vibrio vulnificus*, *Vibrio antiquarius, Vibrio nigripulchritudo, Vibrio parahaemolyticus, Vibrio pelagius,* and the unclassified *Vibrio* strain B1ASS3 (*Vibrio* sp.) were identified to carry similar gene clusters (Fig. 5a). Despite the presence of these diverse *Vibrio* species, *V. cholerae* 7PET strains were the most commonly identified with 37 hits (Fig. 5a), likely reflecting their prominent representation in the NCBI database.

**Figure 5.**
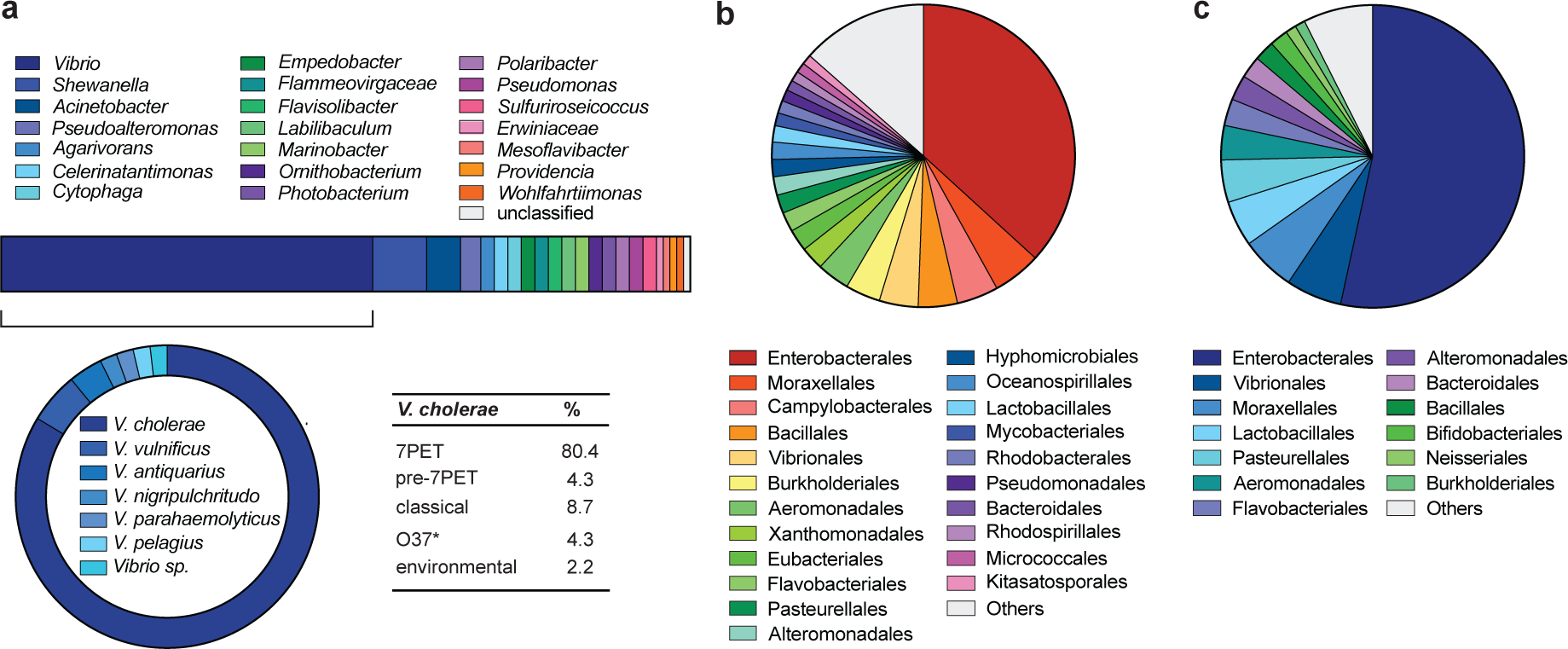
Phylogenetic distribution of the restriction systems. a) The presence of the 5-gene cluster (*VC1765-69*) was assessed across 81,172 bacterial genomes. The results revealed its distribution beyond *V. cholerae*, which represented 54% of all hits. \**V. cholerae* O37 serogroup strains are known to be closely related to classical O1 strains with highly similar chromosomal backbones. **b, c)** Exploration of the (**b**) T1RM system (*VC1769-68-65*) and (**c**) TgvAB system (*VC1766-67*) across the bacterial genomes demonstrates their assessment at the order level of taxonomy. Orders represented in less than 1% of instances were consolidated into a singular category labeled “Others” for the visualization. For details at the species level see Tables S1-S3.

Subsequent analysis focused on the independent occurrences of the T1RM and TgvAB systems. As expected, the T1RM system was widespread (3886 hits) across numerous bacterial orders (Fig. 5b and Table S2 for species-level details). Homologs of the *tgvAB* operon alone were slightly less common with 1744 hits (Table S3 for species-level details), yet 17-times more prevalent than the instances of the five-gene operon described above. Indeed, as shown in Figure 5c, the occurrence of TgvAB homologs spans a wide array of bacterial orders, with species found in the human gut, like *Bacteroides fragilis* (Bacteroidales), to organisms isolated from permafrost, such as *Psychrobacter cryohalolentis* (Moraxellales).

## Conclusion

In this study, we aimed to characterize the predicted restriction gene cluster of VPI-2. We showed that the T1RM system actively methylates the genomes of 7PET *V. cholerae* strains, while restricting unmethylated foreign DNA. Additionally, we identified a novel two-protein modification-dependent restriction system, TgvAB, which is embedded within the T1RM cluster. Interestingly, Picton *et al.* demonstrated that the TgvB homolog BrxU, along with the Bacteriophage Exclusion (BREX) system (55), work in concert to offer complementary resistance against both modified and non-modified phages (44, 48). Therefore, it is tempting to speculate that the embedding of the *tgvAB* operon within the T1RM cluster serves a similar complementary role in *V. cholerae*. Supporting this notion, Machnicka *et al.* noted that GmrSD homologs are frequently encoded within Type I RM loci. An example includes the gene encoding the DUF262 domain-containing protein RloF of *Campylobacter jejuni*, which is situated between *hsdR* and *hsdS* of a T1RM operon (56), similar to the positioning of *tgvAB* described in this study. That defence system tend to cluster together within defence islands has been established over several years (11, 27, 57). However, this concept was recently extended by Payne and colleagues by identifying specific genes embedded within multi-gene defence clusters, highlighting the complex organization and integration of these systems within bacterial genomes (58). Notably, their research found GmrSD-like genes embedded within Hma (Helicase, Methylase, ATPase) defence gene clusters. However, unlike the HEC-05 (=BrxU) and HEC-06 GmrSD-like proteins identified in their work, which function independently (52), our findings indicate that the TgvAB defence operates as a two-protein system, underscoring the diversity in bacterial defence strategies.

## Material and Methods

### Bacterial strains, plasmids, and culture conditions

The bacterial strains and the plasmids used in this study are listed in Table S4. pUC18-mini-Tn7T-Gm-*lacZ* was a gift from Herbert Schweizer via Addgene plasmid #63120 (59). The primary *V. cholerae* strain used, A1552, is a fully sequenced toxigenic O1 El Tor Inaba strain, representing the ongoing 7^th^ cholera pandemic (60, 61). Unless stated otherwise, bacteria were aerobically cultured in Lysogeny broth (LB; 1% tryptone, 0.5% yeast extract, 1% sodium chloride; Carl Roth, Switzerland) with shaking at 180 r.p.m., or on LB agar plates at either 30°C or 37°C. When required, antibiotic selection was applied using ampicillin (100 μg/ml), kanamycin (75 μg/ml), and gentamicin (25 or 50 μg/ml). For natural transformation, chitin powder (Alfa Aesar via Thermo Fisher, USA) was combined with half-concentrated Instant Ocean medium (Aquarium Systems) and sterilized by autoclaving prior to adding the bacterial cultures.

Conjugation with MFDpir (62) was used to introduce the mini-Tn7 transposon derivatives into *E. coli* strain MG1655 on agar plates containing 0.3 mM diaminopimelic acid (DAP; Sigma-Aldrich). To induce expression from the *P*_BAD_ promoter, cultures were grown in media containing 0.2% L-arabinose. For bacteriophages experiments, LB medium was supplemented with 5 mM CaCl_2_ + 20 mM MgSO_4_. Double-layer LB plates were prepared by adding 0.5% agar for semi-solid agar and 1.5% agar for the solid base.

## Genetic engineering of strains and plasmids

Standard molecular cloning techniques were utilized for the cloning process (63) using the following enzymes: Pwo polymerase (Roche), Q5 High fidelity polymerase (New England Biolabs), GoTaq polymerase (Promega), restriction enzymes (New England Biolabs), and T4 DNA ligase (New England Biolabs). Enzymes were used according to the manufacturer’s instructions. All constructs were verified through PCR and/or Sanger or Nanopore sequencing (performed by Microsynth AG, Switzerland) and analysed using SnapGene version 4.3.11.

*V. cholerae* strains were created through natural transformation and FLP recombination (TransFLP) (64–66) or through allelic exchange using derivatives of the suicide plasmid pGP704-Sac28 (67) and SacB-based counter-selection on NaCl-free LB plates with 10% sucrose. Mini-Tn7 transposons, containing *araC* and the gene(s) of interest regulated by the arabinose-inducible promoter *P*_BAD_, were inserted in *E. coli* downstream of *glmS* via triparental mating, following established protocols (68). Site-directed mutations in these constructs were introduced by inverse PCR prior to their transposition into the *E. coli* chromosome.

## PacBio (SMRT) sequencing

Genomic DNA was purified from overnight cultures using Qiagen’s Genomic-tip procedure combined with the Genomic DNA buffer set (Qiagen, Switzerland), following the manufacturer’s instructions. Sample processing, PacBio Single Molecule, Real-Time (SMRT) sequencing, and *de novo* genome assembly were performed at the University of Lausanne’s Genomic Technology Facility, as previously described (35). All SMRT sequencing raw data have been made available on Zenodo (three datasets: 10.5281/zenodo.10838595; 10.5281/zenodo.10839511; 10.5281/zenodo.10839547). Note that the assembled genomes of strains A1552, C6706, C6709, P27459, E7946, DRC193A, and MO10 have been previously reported without analysis of their epigenetic modifications (35, 36, 61).

## Electroporation-mediated transformation of *V. cholerae* using plasmids

To explore the T1RM system’s efficiency in restricting DNA with specific recognition sequences, we compared the uptake frequency of a plasmid harboring the putative recognition motif (P_motif+_) to that of a variant plasmid with silent mutations in *aacC1* (P_motif-_) altering its sequence while maintaining the encoded aminoglycoside-3-*O*-acetyltransferase-I protein. Transformation frequencies were assessed through electroporation. *V. cholerae* competent cells were prepared by standard protocols (63), involving 1:100 dilution of overnight cultures, growth for 2h and 30min at 37°C (OD600∼1.0), and washing steps with cold 2 mM CaCl2 and 10% glycerol before shock-freezing. After 2h at-80°C, electroporation with 100 ng plasmid was performed at 1.6 kV followed by recovery in 2xYT-rich medium at 30 °C for 2 h. Cells were plated on LB agar with and without kanamycin and incubated at 37°C overnight. Transformation frequencies were calculated as the ratio of kanamycin-resistant transformants to the total number of bacteria.

## Bacteriophage handling and culturing

The *E. coli* BASEL phage collection (42) was used in this study. To generate phage stocks, an *E. coli* MG1655ΔaraCBAD (69) overnight culture was diluted and grown to the exponential phase in LB medium supplemented with 5 mM CaCl_2_ and 20 mM MgSO_4_. Subsequently, the culture was 1:10 diluted in prewarmed medium, infected with 10^4^ plaque forming units (PFU)/mL, and incubated under shaking conditions at 37°C for 5 h. Following incubation, centrifugation and filtration were used to clear the lysate, which was then treated with 1% chloroform and stored at +4°C. Phage titers were determined using plaque assays on the propagation strain.

## Bacteriophage plaque assays

For plaque assays, *E. coli* MG1655Δ*araCBAD*, either with the candidate defence system or the empty miniTn7 transposon control, was grown in LB medium. Overnight cultures were diluted 1:100 in LB medium supplemented with 0.2% arabinose, 5 mM CaCl_2_, and 20 mM MgSO_4_ and grown at 37°C with shaking for 2 h. Once reaching the exponential phase, the cultures were diluted 1:40 in 0.5% LB agar containing 5 mM CaCl_2_, 20 mM MgSO_4_, and 0.2% arabinose, then overlaid on 1.5% LB agar. Phage samples were serial diluted in LB medium with 5 mM CaCl_2_ and 20 mM MgSO_4_ and spotted onto the bacterial overlays. After overnight incubation at 37°C, plaques were counted to assess the defense system’s effectiveness compared to the miniTn7-carrying control strain (= fold protection).

## Infection kinetics

The infection kinetics assay of *Tequatrovirus* (Bas35, Bas40) and *Mosigviruses* (Bas46, Bas47) was conducted as follows: Overnight cultures of *E. coli* strains were diluted 1:100 in LB medium supplemented with 5 mM CaCl_2_, 20 mM MgSO_4_ and 0.2% arabinose. Bacterial cultures were then incubated at 37°C with shaking for 2 h. Subsequently, 20 μl of phage per well at multiplicities of infection (MOI) of 0, 0.2, 5, or 10 were added in technical triplicate to a 96-well plate. The cultures were further diluted 1:10 in the same LB condition, and 180 μl of each diluted culture was then added to the wells. The SpectraMax i3x plate reader from Molecular Devices was utilized to assess bacterial growth at 37°C, with measurements taken at 6-minute intervals over a total of 49 cycles. To calculate the MOI, cultures of strains MG1655Δ*araCBAD*-Tn-empty and MG1655Δ*araCBAD*-TnTgvAB were cultured following the protocol outlined in the Bacteriophage Plaque Assays section. Colony-forming units (per ml) were quantified by spotting serially diluted cultures onto LB plates. The calculated values represent the average of three technical replicates.

## Bioinformatics analyses

The VPI-2 genomic region of 7PET O1 strains and one O139 serogroup strain (MO10) was compared and visualized using Clinker software (v 0.0.25, default parameters) (70) after reannotation of the genome sequence of strain A1552 using the Prokaryotic Genome Annotation pipeline version 2023-10-03.build7061 (71) to unify the annotation method. For sequence similarity, NCBI’s blastp was utilized (default parameters, non-redundant protein database; accession August 2023), while structural modeling was conducted with ColabFold (1.5.2) (72) based on AlphaFold2 (51) using default settings. DALI was employed for structural similarity predictions against the Protein Data Bank (PDB) (73, 74).

The distribution of the specific five-gene operon (*VC1765-69*) across bacterial species was examined with MacSyFinder (v.2.1) (75), using a comprehensive database of all sequenced and fully assembled bacterial genomes (taxid:2) from the NCBI database (accession date 26.01.2024). This analysis therefore covered a dataset comprising 81,172 bacterial genomes, which altogether contained over 304 million protein sequences.

To build HMM profiles for each target CDS within the *VC1765-69* operon, homologous protein sequences were identified via PSI-blast searches in the NCBI database (3 iterations), using the non-redundant protein sequence database (accessed in February 2024) with a cutoff e-value of 1e-10.

After identifying homologous sequences for each CDS through PSI-blast, the sequences were aligned using MAFFT (v7.508,--maxiterate 1000 –localpair parameters for higher accuracy alignments) (76). From these multiple alignments, HMM profiles were generated with HMMER (v3.3.2, using hmmbuild with default parameters) (77), forming the basis for constructing different models in MacSyFinder. These models were used to search for the occurrence of the CDS in various combinations encompassing the T1RM and/or TgvAB system genes. The constructed models were then applied in a search across the bacterial protein database mentioned above.

## Statistics and reproducibility

Results are derived from biologically independent experiments, as specified in the figure legends. Statistical analyses were conducted using Prism software (v10.2.1; GraphPad).

## Acknowledgements

We thank members of the Blokesch lab and especially Nicolas Flaugnatti and Alexis Proutière for fruitful discussions and Sandrine Stutzmann, Laurie Righi, and Loriane Bader for technical assistance. We acknowledge the staff of the Lausanne Genomic Technologies Facility and Christian Iseli & Nicolas Guex from the EPFL/UNIL Bioinformatics Competence Center for sample processing, sequencing, genome assembly, and SMRT sequencing analysis. We are also grateful to Alexander Harms for sharing the BASEL phage collection and for valuable discussions. Our appreciation goes to the Waters lab for coordinating the timing of their submission with ours. This work was supported by the Swiss National Science Foundation (310030_185022) and an International Research Scholarship by the Howard Hughes Medical Institute (HHMI) (grant 55008726) awarded to M.B.

## Author contributions

MB secured funding; MB & DWA conceived and supervised the study; GV, DWA, and MB designed the experiments and analysed the data. GV, DWA, and MB designed and constructed strains/plasmids. GV performed the experiments. MB initiated the SMRT sequencing. AL performed comparative genomic and conservation analyses. MB, DWA, and GV wrote the manuscript. All authors approved the final version of the paper.

## Supplemental Material

**Table S1.** Summary of matching hits for the T1RM + TgvAB (*VC1765-69*) model, detected by MacSyFinder v.2.

**Table S2.** Summary of matching hits for the T1RM (*VC1769-68-65*) model, detected by MacSyFinder v.2.

**Table S3.** Summary of matching hits for the TgvAB (*VC1766-67*) model, detected by MacSyFinder v.2.

**Table S4**. Bacterial strains and plasmids used in this study.

## Supplementary Figure

**Figure S1. Observed defence activity against the BASEL phage collection.** Protection levels (fold-protection, as shown by the colour code on the right) were determined by comparing plaque formation in strains with the system to those without, using tenfold serial dilution assays. Data represent the average of two replicates. Details as in Fig. 3a.

