## Supplemental Material for "*Vibrio cholerae* pathogenicity island 2 encodes two distinct types of restriction systems"

### Table of content

**Figure S1.** Observed defence activity against the BASEL phage collection**.**

**Table S1.** Summary of matching hits for the T1RM + TgvAB (*VC1765-69*) model, detected by MacSyFinder v.2.

**Table S2.** Summary of matching hits for the T1RM (*VC1769-68-65*) model, detected by MacSyFinder v.2.

**Table S3.** Summary of matching hits for the TgvAB (*VC1766-67*) model, detected by MacSyFinder v.2.

**Table S4**. Bacterial strains and plasmids used in this study.

**Supplemental References**

### Figure S1.

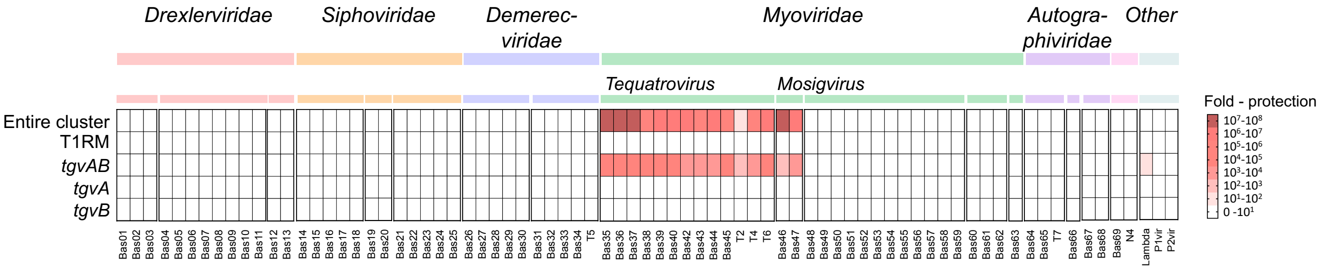

**Figure S1. Observed defence activity against the BASEL phage collection.** Protection levels (fold-protection, as shown by the colour code on the right) were determined by comparing plaque formation in strains with the system to those without, using tenfold serial dilution assays. Data represent the average of two replicates. Details as in Fig. 3a.

**Table S1. Summary of matching hits for the T1RM + TgvAB (*VC1765-69*) model, detected by MacSyFinder v.2.** The table presents the hits obtained for the five-gene cluster model (*VC1765-69*). For each hit, the corresponding species, protein name, and sequence are given. This table is provided as supplementary data file.

**Table S2. Summary of matching hits for the T1RM (*VC1769-68-65*) model, detected by MacSyFinder v.2.** The table presents the hits obtained for the three gene cluster model T1RM (*VC1769-68-65*). For each hit, the corresponding species, protein name, and sequence are given. This table is provided as supplementary data file.

**Table S3. Summary of matching hits for the TgvAB (*VC1766-67*) model, detected by MacSyFinder v.2.** The table presents the hits obtained for the two gene cluster model TgvAB (*VC1766-67*). For each hit, the corresponding species, protein name, and sequence are given. This table is provided as supplementary data file.

**Table S4. Bacterial strains and plasmids used in this study.**

| Strains or plasmids | Genotype/description [a, b] | Strain number | Ref |
|---|---|---|---|
| ***Vibrio cholerae*** | | | |
| A1552 | Wild type; O1 El Tor Inaba; isolated in 1992, Peruvian origin; $Rif^R$ | MB#1 | (1, 2) |
| C6706 | O1 El Tor Inaba; isolated in 1991, Peru; original isolate; non-mutated *luxO*); kind gift from J.J. Mekalanos | MB#4522 | (3) |
| C6709 | O1 El Tor Inaba; isolated in 1991, Peru; $Strep^R$ | MB#1503 | (4) |
| N16961 | O1 El Tor Inaba; isolated in 1975, Bangladesh; *hapR* frame-shifted; $Strep^R$ | MB#2 | (2) |
| P27459 | O1 El Tor Inaba; isolated in 1976, Bangladesh; $Strep^R$ | MB#1504 | (5) |
| E7946 | O1 El Tor Ogawa; isolated in 1978, Bahrain; $Strep^R$ | MB#2600 | (6) |
| DRC193A | O1 El Tor; isolated in 2011, Democratic Republic of Congo; $Strep^R$ | MB#1954 | (7) |
| MO10 | O139 strain isolated in 1992, India; *hapR* mutated (HapR[R12L]); $Strep^R$ | MB#5 | (8) |
| A1552ΔVPI-2 | A1552 deleted for entire VPI-2 through TransFLP; A1552ΔVPI-2::FRT; $Rif^R$ | MB#10090 | This study |
| A1552-VPI-2Δ#2 (Δ*VC1765-1769*) | A1552 with deleted region #2 (Δ*VC1765-1769*::FRT-*kan*-FRT); $Rif^R$ | MB#8678 | (9) |
| A1552Δ*ddmABC* | A1552Δ*ddmABC*-noFRT; *ddmABC* deleted using suicide plasmid pGP704Δ*ddmABC*; $Rif^R$ | MB#9747 | This study |
| A1552Δ*ddm*ABC Δ*VC1765-1769*:: FRT-*kan*-FRT | A1552Δ*ddmABC* deleted for *VC1765-1769* using Δ*VC1765-1769*::FRT-*kan*-FRT PCR fragment; $Kan^R$, $Rif^R$ | MB#11377 | This study |
| A1552Δ*ddmABC* Δ*VC1765-1769* | A1552Δ*ddmABC*Δ*VC1765-1769*::FRT-*kan*-FRT after flip and cure resulting in A1552Δ*ddmABC*Δ*VC1765-1769*::FRT; $Rif^R$ | MB#11379 | This study |
| ***Escherichia coli*** | | | |
| S17-1λ*pir* | TpR SmR *thi pro hsdR- hsdM+ recA* RP4-2-Tc::Mu-Km::Tn7 (λ*pir*) | MB#648 | (10) |
| MFD*pir* | MG1655 RP4-2-Tc::[ΔMu1::*aac(3)IV*-Δ*aphA*- Δ*nic*35-ΔMu2::*zeo*] Δ*dapA*::(*erm-pir*) Δ*recA* | MB#4662 | (11) |
| TOP10 | F- *mcrA* Δ(*mrr-hsdRMS-mcrBC*) Φ80*lacZ*ΔM15 Δ*lac*X74 *recA1 araD*139 Δ(*ara leu*) 7697 *galU galK rpsL* ($Strep^R$) *endA*1 *nupG* | MB#741 | Invitrogen |

| | | | |
|---|---|---|---|
| MFD*pir* / pGP704-Tnempty | MFD*pir* carrying pGP704-Tn*araC* | MB#8793 | This study |
| MFD*pir* / pGP704-Tncluster | MFD*pir* carrying pGP704-TnCluster | MB#11371 | This study |
| MFD*pir* / pGP704-TnT1RM | MFD*pir* carrying pGP704-TnT1RM | MB#11367 | This study |
| MFD*pir* / pGP704-Tn*tgvAB* | MFD*pir* carrying pGP704-Tn*tgvAB* | MB#11368 | This study |
| MFD*pir* / pGP704-Tn*tgvA* | MFD*pir* carrying pGP704-Tn*tgvA* | MB#11369 | This study |
| MFD*pir* / pGP704-Tn*tgvB* | MFD*pir* carrying pGP704-Tn*tgvB* | MB#11370 | This study |
| MG1655Δ*araCBAD* | F- λ-, rph-1 Δ*araCBAD* | MB#4441 | (12) |
| MG1655Δ*araCBAD*-Tnempty | MG1655Δ*araCBAD* with integrated Tn*araC* | MB#10858 | This study |
| MG1655Δ*araCBAD*-TnT1RM | MG1655Δ*araCBAD* with integrated TnT1RM | MB#11372 | This study |
| MG1655Δ*araCBAD*-TnCluster | MG1655Δ*araCBAD* with integrated TnCluster | MB#11373 | This study |
| MG1655Δ*araCBAD*-Tn*tgvAB* | MG1655Δ*araCBAD* with integrated Tn*tgvAB* | MB#11374 | This study |
| MG1655Δ*araCBAD*-Tn*tgvA* | MG1655Δ*araCBAD* with integrated Tn*tgvA* | MB#11375 | This study |
| MG1655Δ*araCBAD*-Tn*tgvB* | MG1655Δ*araCBAD* with integrated Tn*tgvB* | MB#11376 | This study |
| MFD*pir* / pGP704-Tn*tgvA*[G94A]*B* | MFD*pir* carrying pGP704-Tn*tgvA*[G94A]*B* | MB#11382 | This study |
| MFD*pir* / pGP704-Tn*tgvA*[R97A]*B* | MFD*pir* carrying pGP704-Tn*tgvA*[R97A]*B* | MB#11383 | This study |
| MFD*pir* / pGP704-Tn*tgvAB*[G81A] | MFD*pir* carrying pGP704-Tn*tgvAB*[G81A] | MB#11380 | This study |
| MFD*pir* / pGP704-Tn*tgvAB*[R84A] | MFD*pir* carrying pGP704-Tn*tgvAB*[R84A] | MB#11381 | This study |
| MFD*pir* / pGP704-Tn*tgvAB*[H571A] | MFD*pir* carrying pGP704-Tn*tgvAB*[H571A] | MB#11384 | This study |
| MFD*pir* / pGP704-Tn*tgvAB*[N593A] | MFD*pir* carrying pGP704-Tn*tgvAB*[N593A] | MB#11385 | This study |
| MFD*pir* / pGP704-Tn*tgvAB*[N602A] | MFD*pir* carrying pGP704-Tn*tgvAB*[N602A] | MB#11386 | This study |
| MG1655Δ*araCBAD*-Tn*tgvA*[G94A]*B* | MG1655Δ*araCBAD* with integrated Tn*tgvA*[G94A]*B* | MB#11389 | This study |
| MG1655Δ*araCBAD*-Tn*tgvA*[R97A]*B* | MG1655Δ*araCBAD* with integrated Tn*tgvA*[R97A]*B* | MB#11390 | This study |
| MG1655Δ*araCBAD*-Tn*tgvAB*[G81A] | MG1655Δ*araCBAD* with integrated Tn*tgvAB*[G81A] | MB#11387 | This study |

| | | | |
|---|---|---|---|
| MG1655Δ*araCBAD*-Tn*tgvAB*[R84A] | MG1655Δ*araCBAD* with integrated Tn*tgvAB*[R84A] | MB#11388 | This study |
| MG1655Δ*araCBAD*-Tn*tgvAB*[H571A] | MG1655Δ*araCBAD* with integrated Tn*tgvAB*[H571A] | MB#11391 | This study |
| MG1655Δ*araCBAD*-Tn*tgvAB*[N593A] | MG1655Δ*araCBAD* with integrated Tn*tgvAB*[N593A] | MB#11392 | This study |
| MG1655Δ*araCBAD*-Tn*tgvAB*[N602A] | MG1655Δ*araCBAD* with integrated Tn*tgvAB*[N602A] | MB#11393 | This study |
| TOP10 / $P_{[motif +]}$ (pUC-Kan-mTn7T-Gm-*lacZ*) | TOP10 carrying plasmid $P_{[motif +]}$; $Kan^R$; $Gent^R$ | MB#10698 | This study |
| TOP10 / $P_{[motif -]}$ | TOP10 carrying plasmid $P_{[motif -]}$; $Kan^R$; $Gent^R$ | MB#11378 | This study |
| **Plasmids** | | | |
| pBR-FLP | pBR322 derivative containing FLP+, λ cI857+, λ pR from pCP20 integrated into the EcoRV site of pBR322, used for FLP recombination; $Amp^R$ | MB# 1203 | (13) |
| pUX-BF13 | pUX-BF13 - *ori*R6K, helper plasmid with Tn7 transposition function; $Amp^R$ | MB#457 (S17-1λ*pir*) MB#4933 (MFD*pir*) | (14) |
| pGP704-Sac28 | pGP704-Sac28, *ori*R6K, *sacB*; $Amp^R$ | MB#649 | (15) |
| pGP704Δ*ddmABC* | pGP704-Sac28 with Δ*ddmABC*; $Amp^R$ | MB#9727 | This study |
| pGP704-Tn*araC* | pGP704-Tn*araC*; pGP704 with miniTn7 carrying *araC* and the *araBAD* promoter ($P_{BAD}$-); $Amp^R$, $Gent^R$ | MB#5513 (S17-1λpir) MB#8793 (MFD*pir*) | (3, 16) |
| pGP704-TnCluster | pGP704 with miniTn7 carrying *araC* and $P_{BAD}$-*VC1765-69*; $Amp^R$, $Gent^R$ (TnCluster) | MB#11371 | This study |
| pGP704-TnT1RM | pGP704 with miniTn7 carrying *araC* and $P_{BAD}$ -*VC1769-68-65*; $Amp^R$, $Gent^R$ (TnT1RM) | MB#11367 | This study |
| pGP704-Tn*tgvAB* | pGP704 with miniTn7 carrying *araC* and $P_{BAD}$ -*VC1766-67*; $Amp^R$, $Gent^R$ (Tn*tgvAB*) | MB#11368 | This study |
| pGP704-Tn*tgvA* | pGP704 with miniTn7 carrying *araC* and $P_{BAD}$ -*VC1767*; $Amp^R$, $Gent^R$ (Tn*tgvA*) | MB#11369 | This study |
| pGP704-Tn*tgvB* | pGP704 with miniTn7 carrying *araC* and $P_{BAD}$ -*VC1766*; $Amp^R$, $Gent^R$ (Tn*tgvB*) | MB#11370 | This study |
| pGP704-Tn*tgvA*[G94A]*B* | pGP704 with miniTn7 carrying *araC* and $P_{BAD}$ -*VC1766-67* encoding G94A variant of *VC1767*; $Amp^R$, $Gent^R$ (Tn*tgvA*[G94A]B) | MB#11382 | This study |

| | | | |
|---|---|---|---|
| pGP704-Tn*tgvA*[R97A]*B* | pGP704 with miniTn7 carrying *araC* and $P_{BAD}$ -*VC1766-67* encoding R97A variant of *VC1767*; Amp$^R$, Gent$^R$ (Tn*tgvA*[R84A]B) | MB#11383 | This study |
| pGP704-Tn*tgvAB*[G81A] | pGP704 with miniTn7 carrying *araC* and $P_{BAD}$ -*VC1766-67* encoding G81A variant of *VC1766*; Amp$^R$, Gent$^R$ (Tn*tgvAB*[G81A]) | MB#11380 | This study |
| pGP704-Tn*tgvAB*[R84A] | pGP704 with miniTn7 carrying *araC* and $P_{BAD}$ -*VC1766-67* encoding R84A variant of *VC1766*; Amp$^R$, Gent$^R$ (Tn*tgvAB*[R84A]) | MB#11381 | This study |
| pGP704-Tn*tgvAB*[H571A] | pGP704 with miniTn7 carrying *araC* and $P_{BAD}$ -*VC1766-67* encoding H571A variant of *VC1766*; Amp$^R$, Gent$^R$ (Tn*tgvAB*[H571A]) | MB#11384 | This study |
| pGP704-Tn*tgvAB*[N593A] | pGP704 with miniTn7 carrying *araC* and $P_{BAD}$ -*VC1766-67* encoding N593A variant of *VC1766*; Amp$^R$, Gent$^R$ (Tn*tgvAB*[N593A]) | MB#11385 | This study |
| pGP704-Tn*tgvAB*[N602A] | pGP704 with miniTn7 carrying *araC* and $P_{BAD}$ -*VC1766-67* encoding N602A variant of *VC1766*; Amp$^R$, Gent$^R$ (Tn*tgvAB*[N602A]) | MB#11386 | This study |
| pUC18-miniTn7T-Gm-*lacZ* | pUC18-miniTn7T-Gm-*lacZ*; Amp$^R$, Gent$^R$ | MB#10656 | Addgene # 63120 (17) |
| $P_{[motif +]}$ | pUC-Kan-mTn7T-Gm-*lacZ* derived from pUC18-miniTn7T-Gm-*lacZ* by replacing *bla* (Amp$^R$) by *aph* (Kan$^R$) cassette; harboring motif [GATGCCCATACTT] within *aacC1* (Gent$^R$); Kan$^R$, Gent$^R$ | MB#10698 | This study |
| $P_{[motif -]}$ | pUC-Kan-mTn7T-Gm-*lacZ*, harboring silent mutations within *aacC1* (Gent$^R$) to modify motif [**A**AT**A**CCCAT**TGAA**]; Kan$^R$, Gent$^R$ | MB#11378 | This study |

[a] *V. cholerae* locus tag numbers are according to Heidelberg *et al.* (18)
[b] Kan – Kanamycin; Rif – Rifampicin; Strep – Streptomycin; Gent – Gentamicin; Amp – Ampicillin
